# Bridging biology and statistics with hybrid Bayesian experimental design for drug dose-response assays

**DOI:** 10.1101/2025.09.18.677072

**Authors:** Atlanta Chakraborty, Siyi Chen, Xun Huan, Gary Luker

## Abstract

Dose-response cytotoxicity assays are central to evaluating drug potency, yet selecting concentrations that capture both the full response range and key parameters such as the half-maximal inhibitory concentration (IC_50_) and Hill slope remains challenging. Practical constraints, including limited replicates, variability across culture conditions, and the trial-and-error nature of current practice, make assays time- and resource-intensive. To address this, we introduce a Bayesian inference framework that quantifies uncertainty, incorporates prior knowledge, and extend it with a Bayesian optimal experimental design (OED) strategy to systematically refine concentration selection. We further propose a hybrid OED approach that integrates information-theoretic design with space-filling principles, aligning statistical rigor with biological intuition. Applied to ER+ breast cancer cells in monoculture and co-culture with stromal cells, this framework revealed differential drug responses while improving information efficiency. More broadly, our results highlight how Bayesian OED can bridge computational and biological perspectives, offering a path toward more efficient, reproducible, and interpretable experimental design in cancer research and beyond.

**Author summary:** We present a statistical framework that systematically captures uncertainty in drug cytotoxicity assays, which are standard laboratory tests that measure how sensitive cells are to drug treatments. Applying this approach to breast cancer cells grown with and without supportive bone marrow cells, we revealed how the surrounding environment can protect cancer cells from therapy. By guiding the selection of drug concentrations, our framework reduces trial-and-error, thereby saving time and resources. Beyond cancer research, this strategy offers a general way to design more efficient biological experiments, ultimately supporting the development of new treatments.

## 1 Introduction

### 1.1 Drug dose-response assays

Cell viability assays, often referred to as cytotoxicity assays, are a common technique in cell culture for quantifying cell survival following exposure to stressors such as chemotherapy drugs. Readouts of cell viability include metabolic activity, adenosine triphosphate (ATP) content, or total protein levels, among others. These assays are widely used for applications ranging from optimizing cell culture conditions and measuring cell proliferation to assessing survival under various treatments [1].

In cancer research, a two-dimensional (2D) cell-based assay, known as a *drug dose-response assay*, is frequently employed to evaluate the potency of therapeutic agents under diverse conditions [2]. In our previous work, we used this approach to examine drug resistance in estrogen receptor-positive (ER+) breast cancer cells co-cultured with mesenchymal stem cells (MSCs) [3]. By comparing the viability of cancer cells cultured alone versus co-cultured across a range of drug concentrations, we gained insights into how stromal cells in the tumor environment regulate resistance to therapy.

Conducting such assays, however, is laborious and resource-intensive. Cells must be cultured under precise conditions, seeded into multi-well plates, treated with carefully pipetted drug concentrations, and incubated before viability measurements are collected. These measurements are then used to estimate key parameters of the dose-response curve, including the half-maximal inhibitory concentration (IC_50_) and the Hill slope (HS). Selecting drug concentrations that reliably reveal these parameters is nontrivial. Experimenters often rely on evenly spaced concentrations, refining them through multiple rounds of testing. This trial-and-error process is time-consuming, costly, and can limit reproducibility across laboratories.

### 1.2 Bayesian uncertainty quantification

To address these challenges, we adopt a Bayesian framework that represents uncertainty explicitly and updates it systematically as new data become available [4–9]. Unlike the frequentist view of probability as long-run frequency, the Bayesian perspective interprets probability as a degree of belief. Unknown model parameters denoted by *θ* (such as IC_50_ and HS) are represented as random variables with an associated *prior distribution p*(*θ*), reflecting initial uncertainty before data are observed.

When an experiment is performed at a design condition *ξ* (e.g., a chosen drug concentration) and produces measurement data *y*, the prior distribution is updated via Bayes’ rule:

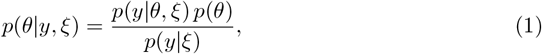

where *p*(*θ* | *y, ξ*) is the *posterior distribution* representing updated knowledge after incorporating the data; *p*(*y* | *θ, ξ*) is the *likelihood* of observing *y* given parameters *θ* under design *ξ*; and *p*(*y* | *ξ*) is the *marginal likelihood* that normalizes the distribution.

Bayesian updating provides a principled way to aggregate evidence across sequential experiments while avoiding under- or over-confidence. It is especially advantageous in biology [10–12], where measurements are often sparse, noisy, or indirect, and where incorporating prior knowledge from historical data, expert judgment, or related systems can accelerate inference.

### 1.3 Experimental design strategies

Experiments and data collection are fundamental to scientific research, but they are often delicate, costly, and time-consuming. A systematic approach to quantifying and optimizing the value of experiments is therefore essential. Two broad traditions of experimental design have emerged.

#### Design of experiments (DOE)

DOE [13, 14] emphasizes exploration without requiring a predictive model of the system. These morel-free approaches are intuitive, robust, and widely used by experimentalists. They include space-filling designs that distribute points broadly across a domain, such as maximin designs that maximize the minimum distance between points. Popular techniques include Latin hypercube [15] and maximum projection (MaxPro) designs [16, 17], which seek space-filling properties not only in the full design space but also in lower-dimensional projections. Other approaches aim for uniform point distributions [18]. Standard strategies such as factorial and composite designs [19] [20, Chapter 7] also fall in this category. These methods align with the experimental intuition that “covering the space” provides security against missing important responses.

#### Optimal experimental design (OED)

In contrast, OED [21–25] is model-based and exploitative: it leverages predictive models to simulate potential outcomes and prioritize experiments expected to yield the most information. Bayesian OED further builds on the Bayesian framework introduced earlier in Section 1.2, defining a design criterion called the *expected utility U* (*ξ*) to quantify the value of an experiment under design *ξ*. A common choice is the *expected information gain* (EIG), also known as mutual information [26]:

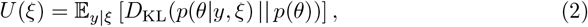

where *D*_KL_ is the Kullback–Leibler (KL) divergence from the prior *p*(*θ*) to the posterior *p*(*θ* | *y, ξ*). A larger KL divergence indicates that an experiment substantially reduces parameter uncertainty, making it more informative. The goal is then to find the optimal design:

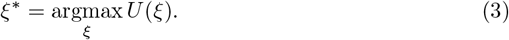

### 1.4 A bridging perspective

While DOE and OED differ in philosophy—coverage versus efficiency—both have merit. DOE resonates with experimental intuition and emphasizes robustness, while OED leverages models to maximize the information yield from limited resources. Our perspective is that these approaches are not in opposition but are complementary.

In this work, we develop a hybrid design strategy that combines the interpretability and robustness of space-filling approaches with the efficiency of Bayesian information-theoretic design. Our goal is to present a framework that is accessible to biologists while demonstrating how statistical models can guide experimental design more systematically. In doing so, we aim to build common ground between experimental practice and statistical theory, advancing both biological discovery and computational methodology.

## 2 Results

### 2.1 Pilot assays and baseline modeling

We began with pilot dose-response assays over a broad concentration range (0–3000 nM) of alpelisib, a phosphatidylinositol 3-kinase inhibitor, using two replicates each for MCF7 cells in monoculture and in co-culture with HS5 stromal cells. The viability data were analyzed using a standard four-parameter logistic (sigmoid) inhibition curve (see Section 4.3 for details):

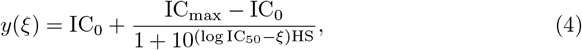

where *y* is the response activity (normalized photon flux), *ξ* is the drug concentration being designed, IC_max_ is the response at the maximum drug concentration, IC_0_ is the baseline response, IC_50_ is the concentration that reduces viability by half, and HS is the Hill slope controlling curve steepness.

Fitting this model to the pilot data suggested systematic differences between monoculture and co-culture (see Figure 1). Co-culture shifted the inhibition curve toward higher IC_50_, consistent with increased drug resistance, and altered the slope, reflecting different response dynamics.

**Fig 1.**
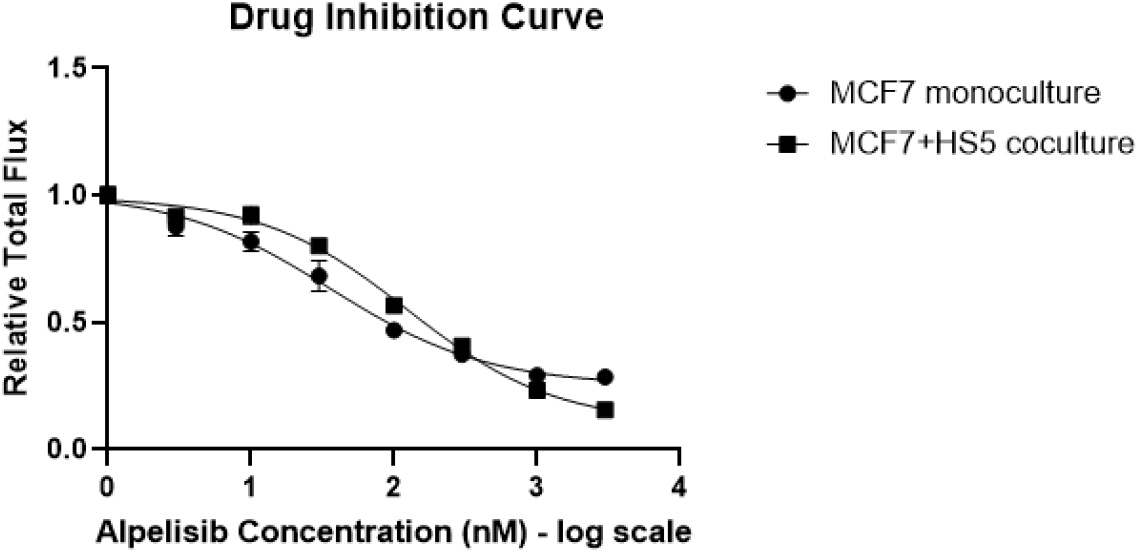
Dose-response curves of MCF7 cells treated with alpelisib under different culture conditions. Monocultures of MCF7 cells or co-cultures of MCF7 with HS5 stromal cells were treated for three days with increasing concentrations of alpelisib. Viability of MCF7 cells was quantified by bioluminescence from stably expressed click beetle luciferase, which is absent in HS5 cells. Values were normalized to vehicle-treated controls (each data point represents quadruplicates). Fitted dose-response curves revealed IC_50_ values of 36.3 and 55.0 (two replicates) for monoculture and 126 and 97.7 nM for co-culture. Corresponding HS values were −0.79 and −0.85 for monoculture, and −0.61 and −0.97 for co-culture.

To rigorously quantify uncertainty in these parameters, we employed a Bayesian hierarchical model similar to that of [10]. The key unknown parameters (IC_50_, HS) were treated as random variables with biologically informed priors, while the likelihood was given by the logistic model in Equation (4) with log-normal measurement noise (see Section 4.3 for details). Posterior distributions (Figure 2) provided a full probabilistic characterization of parameter uncertainty. Their peaks aligned well with point estimate results in Figure 1, but the posterior framework additionally revealed credible regions for the parameters. For example, in co-culture the posterior distributions for both IC_50_ and HS showed significant shifts despite residual uncertainty. These probabilistic results offered a more nuanced view of drug-response heterogeneity, providing transparency on which parameter values are more or less plausible, and laying a strong foundation for subsequent experimental design.

**Fig 2.**
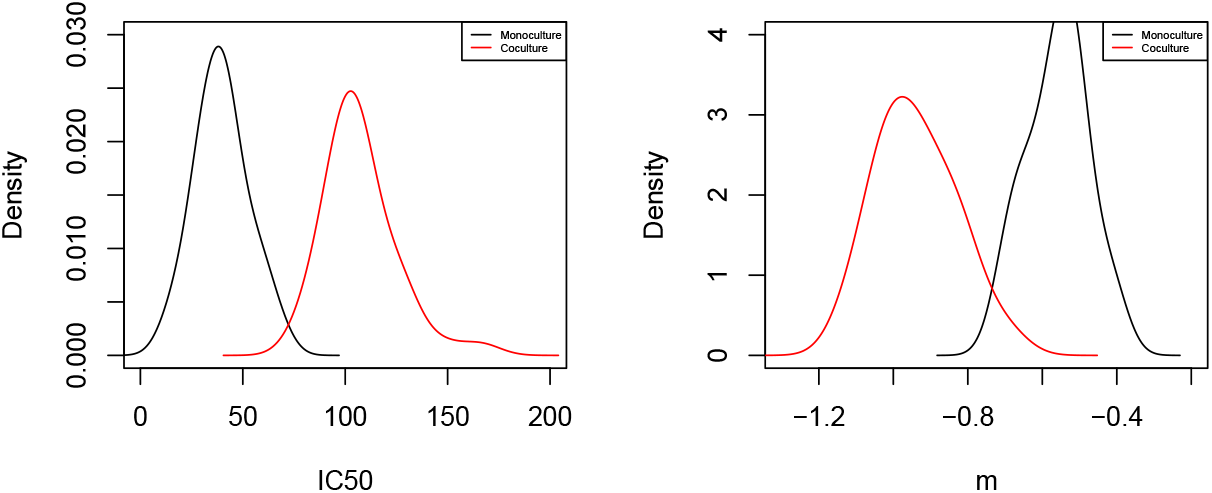
Marginal posterior distributions for monoculture and co-culture conditions. Posterior peaks closely align with the best-fit values from Figure 1, while the probability distributions additionally quantify parameter uncertainty.

### 2.2 First attempt at optimal experimental design

With these baseline results, we next asked whether OED could guide the choice of more informative drug concentrations. Using EIG (Equation (2)) as the design criterion, we identified concentration sets predicted to maximally reduce uncertainty in IC_50_ and HS.

The first OED-generated designs (Table 1) tended to place concentrations near quarter intervals of the dose range when expressed on a log scale. Statistically, this clustering reflects optimality: responses near the steeper regions of the curve provide the most information about slope and inflection. Biologically, however, such designs were less intuitive, as they offered limited coverage of intermediate responses that are important for interpreting cellular behavior. This revealed a key tension: designs that are mathematically helpful are not always aligned with biological practice. Importantly, this mismatch was not a shortcoming of OED but a valuable diagnostic, underscoring the gap between purely information-theoretic design and the experimental intuition of space-filling coverage.

**Table 1.** Concentration values and their log-transformed counterparts from the first OED design. The initial design shows clustering of several points.

| Concentration (nM) | Log-concentration ($\xi$) |
| --- | --- |
| 11 | 1.04 |
| 18 | 1.26 |
| 20 | 1.30 |
| 21 | 1.32 |
| 639 | 2.81 |
| 937 | 2.97 |
| 1268 | 3.10 |
| 1396 | 3.15 |

### 2.3 Hybrid design: integrating information-theoretic and space-filling principles

To reconcile these perspectives, we developed a hybrid design strategy that balances the efficiency of Bayesian OED with the robustness and interpretability of space-filling designs. The hybrid utility function augments EIG with a penalty for closely clustered points, thereby encouraging designs that are both informativeness and well distributed across the dose range:

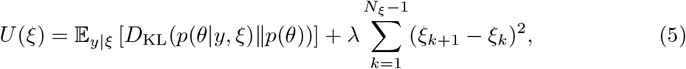

where *ξ* values are expressed in the log-concentration scale to promote spacing in log space, the second term penalizes uneven spacing, and λ *>* 0 controls the trade-off between informativeness and coverage. This balance can be tuned to reflect user priorities or experimental context. To ensure feasibility, all concentrations were rounded to practical pipetting values, and boundary doses (0 and 3000 nM) were always included.

The resulting designs yielded inhibition curves that were both biologically interpretable and statistically informative. Posterior estimates of IC_50_ and HS under the hybrid design captured clear differences between monoculture and co-culture while requiring fewer replicates. For example, a six-dose purely space-filling design achieved an EIG of only 0.24, whereas our hybrid design reached 7.64 while still maintaining good coverage. We then implemented an *N*_*d*_ = 8 hybrid design (Table 2), yielding fitted response curves in Figure 3. This set provided well-distributed sampling across the log scale while delivering substantial information gain.

**Table 2.** Concentration values and their log-transformed counterparts from the hybrid OED design. End-point concentrations of 0 and 3000 nM were always included. Compared with the initial OED design, the selected doses are more evenly distributed across the range, avoiding excessive clustering.

| Concentration (nM) | Log-concentration ($\xi$) |
| --- | --- |
| 0 | 0.00 |
| 147 | 2.17 |
| 340 | 2.53 |
| 488 | 2.69 |
| 554 | 2.74 |
| 738 | 2.87 |
| 1810 | 3.26 |
| 3000 | 3.48 |

**Fig 3.**
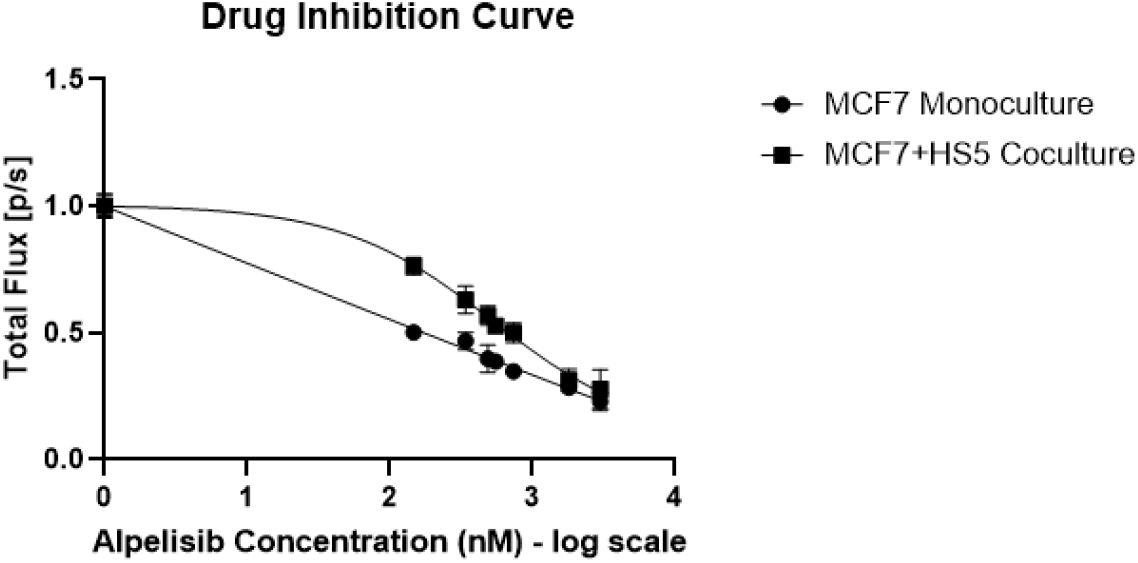
Dose-response curves under the hybrid OED design. With the space-filling penalty added, the selected concentrations were more evenly distributed, reducing redundancy compared with the initial OED design. Using these concentrations, MCF7 monocultures and MCF7–HS5 co-cultures were treated with alpelisib for three days under the same experimental setup. Viability of MCF7 cells was quantified by bioluminescence from stably expressed click beetle luciferase, which is absent in HS5 cells. Values were normalized to vehicle-treated controls (each data point represents quadruplicates). Fitted dose-response curves yielded IC_50_ values of 0.000384 for monoculture and 531 for co-culture, with HS values of −0.032 and −0.82, respectively.

### 2.4 Summary of results

These results demonstrate that a hybrid design strategy can bridge two traditions of experimental practice. Space-filling designs provide robustness but may be inefficient, while pure OED maximizes informativeness but risks impractical clustering. By integrating the two, the hybrid approach respects biological intuition while leveraging statistical models to improve efficiency.

This bridging perspective illustrates the broader potential of Bayesian design in biological research: rather than replacing existing practices, it can complement them to reduce resource use and sharpen biological insight.

## 3 Discussion

In this study, we introduced a Bayesian inference framework for drug dose-response assays and demonstrated its utility in optimizing the selection of drug concentrations. Using ER+ breast cancer cells in monoculture and in co-culture with stromal HS5 cells, we showed that Bayesian modeling captures uncertainty in parameters such as IC_50_ and HS, providing richer insights than point estimates alone. Building on these results, we implemented an OED strategy. While initial designs clustered doses at certain locations, the introduction of a hybrid criterion, combining EIG with a space-filling penalty, yielded concentration sets that were both biologically interpretable and statistically informative.

Traditional cell viability assays typically rely on evenly spaced concentrations with iterative adjustments based on observed responses. This approach is intuitive and robust but often resource-intensive, requiring multiple rounds to refine parameter estimates [2]. The Bayesian framework complements this practice by explicitly quantifying uncertainty and using it to guide design choices. Our hybrid OED strategy preserves the interpretability of evenly spread designs while prioritizing concentrations that maximize EIG. In this way, it integrates experimental intuition with statistical efficiency rather than replacing one with the other.

This ability to systematically identify informative concentrations offers several advantages for experimental biology. First, it reduces the number of replicates and iterative rounds required, thereby conserving both time and resources. Second, it promotes standardization and reproducibility by mitigating variability introduced through *ad hoc* choices of concentration ranges. Third, the Bayesian framework uniquely enables the incorporation of prior knowledge—whether from historical datasets, related experiments, or expert judgment—and provides a probabilistic representation that is especially valuable when data are sparse or noisy. Importantly, its goal is not only to generate parameter estimates but also to honestly represent uncertainty and knowledge gaps, thereby fostering more transparent inference.

Although demonstrated here in the context of alpelisib dose-response assays, the principles of Bayesian OED are broadly generalizable. Similar approaches could be applied to other drugs, cell lines, or experimental systems where resource constraints and uncertainty complicate design. The hybrid framework is flexible: the information-theoretic component can be adapted to target specific quantities of interest, while the space-filling component maintains biological interpretability. This adaptability extends beyond cancer biology to any setting where dose-response or stimulus-response relationships are studied.

At the same time, limitations remain. Bayesian OED can be computationally intensive, potentially creating barriers for laboratories without statistical expertise or resources. Future work should address these barriers through user-friendly software and adaptive design methods that update concentration choices in real time. Like any model-based approach, OED depends on the assumptions of the underlying model, and model misspecification remains a key challenge. While prior distributions enable incorporation of prior knowledge, they also require careful specification and can impose a burden on users. Addressing model error and improving prior construction workflow are important areas for further research.

Looking ahead, several directions are promising. One is to extend the model to more complex experimental settings, such as drug combinations or variable environmental conditions. Another is to improve performance with fewer concentrations and replicates, making the approach more suitable for early-stage studies with limited data. Ultimately, tailoring OED objectives to experimental goals will be crucial: while Equation (2) targets parameter learning, in practice the goals often include prediction, therapy optimization, or decision support. Goal-oriented OED frameworks [27, 28] could align the mathematical design criteria more closely with biological and clinical objectives. Similarly, sequential OED approaches [29–32] that adapt across multiple rounds of experimentation offer opportunities to plan beyond individual assays.

Through Bayesian OED, we presented a framework that bridges computational and biological perspectives on assay design. The hybrid design approach respects experimental intuition while leveraging statistical models to maximize information yield. In doing so, it creates opportunities for more efficient experiments, and contributes to building common ground where statisticians and biologists can collaborate more effectively.

## 4 Materials and methods

### 4.1 Experimental setup for dose-response assay

We used human MCF7 breast cancer cells (ATCC, Manassas, VA, USA) and HS5 immortalized bone marrow mesenchymal stem cells (ATCC). Both cell lines stably expressed click beetle green (CBG) or click beetle red (CBR) luciferases [33]. Cells were cultured in DMEM (#11995, Gibco, Thermo Fisher, Grand Island, NY, USA) supplemented with 10% fetal bovine serum (FBS) (HyClone, ThermoScientific, Waltham, MA, USA), 1% GlutaMAX (#35050, Gibco), 1% Penicillin-Streptomycin (P/S, #15140, Gibco), and 50mg/L Plasmocin™ prophylactic (Invivogen, San Diego, CA, USA).

For viability assays, cells were plated in black-walled, flat-bottom 96-well plates (ThermoFisher Scientific, Waltham, MA, USA). Monocultures consisted of MCF7 alone, while co-cultures contained MCF7:HS5 cells at a 1:9 ratio. A total of 10,000 cells were seeded per well, with quadruplicates per condition. After one day, standard growth medium were changed to low-glucose, low-serum (LG/LF) medium using phenol-red free DMEM (#A14430–01, Gibco) supplemented with 1% FBS, 1% P/S, 1% Glutamax, 1% Sodium Pyruvate, 10 nM-estradiol (#E2758, Sigma-Aldrich, Millipore Sigma, Saint Louis, MO, USA), and 5 mM glucose. Cells were then incubated for an additional 72 hours with various concentrations of alpelisib (#S2814, Selleckchem) [34, 35]. Viability was quantified via bioluminescence imaging of luciferase activity, normalized to vehicle controls.

### 4.2 Data Acquisition

To quantify changes in relative numbers of viable breast cancer and stromal cells in co-cultures, we performed dual-color bioluminescence imaging of CBG (cancer cells) or CBR (stromal cells) using an IVIS Lumina Series III (Perkin Elmer, Waltham, MA, USA) as previously described in [36]. Both CBG and CBR are ATP-dependent luciferase enzymes, so bioluminescence measures relative numbers of metabolically active cells. We exported photon flux data from regions of interest defined in Living Image 4.3.1 (Perkin Elmer) and calculated the mean and standard error of the mean (SEM) for every group of quadruplicates. We expressed data as percent change in bioluminescence relative to vehicle control, calculated as:

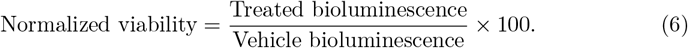

Similarly, for the SEM, we divided the original standard error value by the percent change value in the same group.

### 4.3 Logistic dose-response model and Bayesian setup

We modeled dose-response data with a four-parameter logistic function:

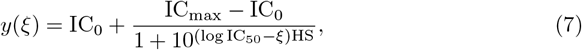

where *y* is the response activity (normalized photon flux), *ξ* is the drug concentration being designed, IC_max_ is the response at the maximum drug concentration, IC_0_ is the baseline response, IC_50_ is the concentration at which viability is reduced by half, and HS is the Hill slope that controls the steepness of the response.

To quantify parameter uncertainty, we adopted a Bayesian hierarchical model following [10]. Each observation followed:

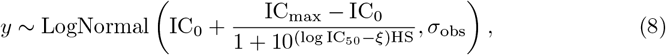

with prior distributions (same for monoculture and co-culture):

- *σ*_obs_ ∼ Cauchy(0, 5);
- IC_0_ ∼ Uniform(0, 0.2);
- IC_max_ ∼ Uniform(0.8, 1);
- IC_50_ ∼ LogNormal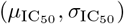 with 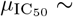 Normal(0, 10) and 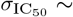 Cauchy(0, 5); and
- HS ∼ LogNormal(*µ*_HS_, *σ*_HS_) with *µ*_HS_ ∼ Normal(0, 10) and *σ*_HS_ ∼ Cauchy(0, 5).

Posterior inference was performed using Markov chain Monte Carlo (MCMC) [37, 38], implemented in R.

Posterior distributions from pilot assays were then projected into simplified uniform priors for subsequent OED. This step prevents overconfidence from limited pilot data, ensuring robustness and generalizability. The updated OED priors for monoculture are:

- *σ*_obs_ ∼ Uniform(0.04, 0.1);
- IC_0_ ∼ Uniform(0.03, 0.2);
- IC_max_ ∼ Uniform(0.95, 1.09);
- IC_50_ ∼ Uniform(26, 107); and
- HS ∼ Uniform(−0.82, −0.42),

and for co-culture are:

- *σ*_obs_ ∼ Uniform(0.05, 0.08);
- IC_0_ ∼ Uniform(0.03, 0.17);
- IC_max_ ∼ Uniform(0.94, 1.02);
- IC_50_ ∼ Uniform(84, 163); and
- HS ∼ Uniform(−1.25, −0.72).

### 4.4 Optimal experimental design

As described in Section 1.3, Bayesian OED was formulated by maximizing the EIG:

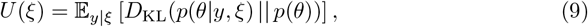

with *ξ* representing dose log-concentrations. To reconcile informativeness with biological interpretability, we introduced a hybrid criterion:

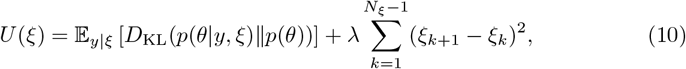

where the second term penalizes clustering, and λ *>* 0 tunes the balance between information gain and spread.

Numerical evaluation of *U* (*ξ*) used a nested Monte Carlo estimator [39, 40]:

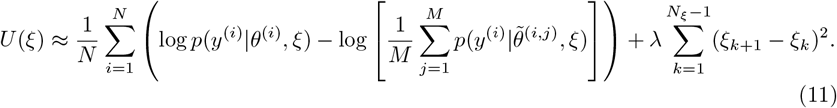

Here, *θ*^(*i*)^ and *θ*^(*i,j*)^ are prior samples, and *y*^(*i*)^ are synthetic datasets simulated from the likelihood.

The design space was defined as [0, 3000] nM on a log scale, with 0 and 3000 nM end points always included. Optimization of *ξ*^∗^ = argmax^*ξ*^ *U* (*ξ*) was performed using Bayesian optimization [41, 42].

## Author Contributions

**Conceptualization:** Xun Huan, Gary Luker

**Formal analysis:** Atlanta Chakraborty, Siyi Chen

**Funding acquisition:** Xun Huan, Gary Luker

**Investigation:** Atlanta Chakraborty, Siyi Chen

**Software:** Atlanta Chakraborty, Siyi Chen

**Supervision:** Xun Huan, Gary Luker

**Visualization:** Atlanta Chakraborty, Siyi Chen

**Writing – original draft:** Atlanta Chakraborty, Siyi Chen

**Writing – review & editing:** Xun Huan, Gary Luker

## Acknowledgments

The authors acknowledge support from United States National Institute of Health grants R01CA238042, R01CA238023, R33CA225549, U24CA237683, R37CA222563, and P30 CA047904. We also acknowledge support from the W. M. Keck Foundation.

